# PreyTouch: An Automated System for Prey Capture Experiments Using a Touch Screen

**DOI:** 10.1101/2024.06.16.599188

**Authors:** Regev Eyal, Mark Shein-Idelson

## Abstract

The ability to catch prey is crucial for survival and reproduction and is subject to strong natural selection across predators. In many animals, prey capture demands the orchestrated activation of multiple brain regions, demonstrating the intricate interplay between sensory processing, decision-making, and motor execution. This makes prey capture a prime paradigm in neuroscience. Further, its ubiquity across species makes it ideal for comparative research and for studying the evolution of cognition. However, despite recent technological advances in the collection and analysis of behavioral data, experimental approaches for studying prey catch are lagging behind. To bridge this gap, we created PreyTouch - a novel system for performing prey capture experiments on a touch screen. PreyTouch incorporates flexible presentation of prey stimulus, accurate monitoring of predator strikes and automated rewarding. The system’s real time processing enables closing the loop between predator movement and prey dynamics for studying predator-prey interactions. Further, the system is optimized for automated long-term experiments and features a web-ui for remote control and monitoring. We successfully validated PreyTouch by conducting long-term prey capture experiments on the lizard *Pogona Vitticeps*. The acquired data revealed the existence of prey preferences, complex prey attack patterns, and fast learning of prey dynamics. The unique properties offered by PreyTouch combined with the ubiquity of prey capture behaviors across animals establish it as a valuable platform for comparatively studying animal cognition.

## Introduction

The ability to successfully identify and capture moving prey is essential for the survival, maturation, and reproduction in many animals. Capturing prey demands substantial time and resources for its planning and successful execution (Abrams, 1984). Consequently, it is a trait under intense selective pressure and plays a pivotal role in shaping evolutionary trajectories (Stephens and Krebs, 1986). Reflecting its importance, prey capture was extensively studied across various disciplines ranging from ecology to psychology (Stephens et al., 2007) and over diverse species spanning multiple animal classes (Ben-Tov et al., 2015; Bianco et al., 2011; Borghuis and Leonardo, 2015; Catania, 2015; Chang et al., 2018; Espinasa et al., 2014; Ewert et al., 2001; Hedenström and Rosén, 2001; Hoy et al., 2016; Jiun-Shian Wu and Chiao, 2023; Konishi, 1973; Mearns et al., 2020; Meyers and Herrel, 2005; Mischiati et al., 2015; Villanueva et al., 2017).

Prey capture, in its complexity, demands an orchestrated activation of multiple brain regions (Ewert et al., 2001; Geberl et al., 2015), showcasing the intricate interplay between sensory processing (Bianco et al., 2011), decision-making (Wang et al., 2023), and motor execution (Mearns et al., 2020). To successfully catch prey, animals need to first segment, identify, localize and sensorily track the prey in space (Mearns et al., 2020). Next, they need to decide whether to attack the prey by weighting multiple available options while taking into account possible risks in their surrounding environment (Pyke et al., 1977). Finally, they need to plan the attack (Ngo et al., 2022) and the strike, requiring the execution of appropriate and precise motor commands (Mearns et al., 2020). Such multifaceted processing is bound to strongly depend on (and drive the evolution of) multiple computations within brain circuits. This, together with the abundance of prey catch behavior across animal classes, makes prey capture a prime paradigm for comparative studies, particularly in the context of evolution of cognition and neural computations (Cisek, 2019; Dukas, 2004). However, to exploit the potential in prey catch investigations, systematic behavioral experiments combined with electrophysiological measurements are needed.

In recent years, the field of animal behavior has undergone a significant transformation with the integration of advanced technologies (Mathis et al., 2018; Robson and Li, 2022; Wiltschko et al., 2015). These technologies enable the collection and analysis of extensive datasets, facilitating the exploration of new questions (Miller et al., 2022) previously inaccessible in classical prey capture studies (Ewert, 1987). A number of pioneering studies have already begun to leverage these technologies to explore some aspects of prey capture (Bianco et al., 2011; Catania, 2015; Michaiel et al., 2020; Mischiati et al., 2015; Szopa-Comley and Ioannou, 2022). However, there remains a lack of open-source platforms for facilitating generalized and automated prey capture experiments, especially for freely moving animals. In particular, one potentially promising technology that has yet to be adopted for systematically studying prey capture in the lab is the use of touch screens (Horner et al., 2013; Kangas and Bergman, 2017; Lewis et al., 2020; Mueller-Paul et al., 2014; Nieder, 2018; Oh et al., 2019).

To bridge this gap, we present PreyTouch - a novel touch screen based system for prey capture experiments. The system incorporates a touch screen application (Figure 1A), for presenting naturalistic and artificial prey dynamics. It supports real-time strike detection and analysis (Figure 1B) as well as routines for feeding this information back to alter prey movement (Figure 1A) and facilitate predator-prey interactions, or to automatically deliver rewards (including live rewards) and reinforce specific choices (Figure 1C). PreyTouch incorporates comprehensive camera control and calibration (Figure 1D) enabling precise frame-by-frame real-time analysis using deep models (Figure 1E). PreyTouch includes a user-friendly web-based interface for remote control, customization, scheduling and monitoring of experiments (Figure 1F) to support automated long-term experiments. PreyTouch ensures high precision synchronization between the system’s components and with external devices (e.g. electrophysiological recordings) (Figure 1G). Finally, all information collected by PreyTouch is logged and can be displayed with viewers for tracking of animal performance (Figure 1H). We validated the system by successfully performing prey catch experiments in the lizard *Pogona Vitticeps*.

**Figure 1:**
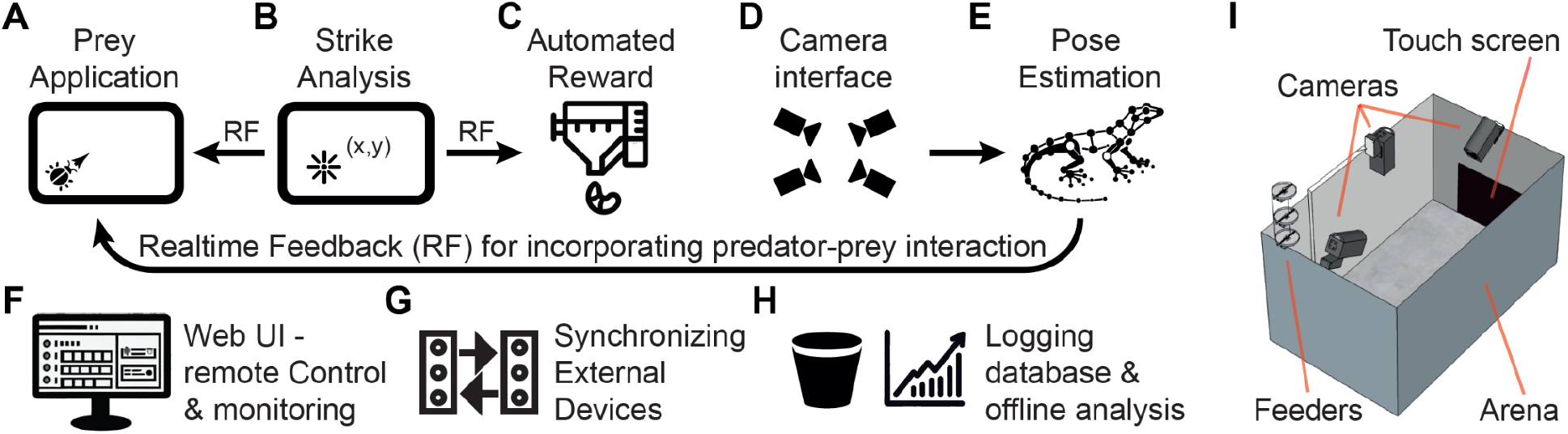
Components of the PreyTouch System. (A) Touch screen prey application allowing flexible animation of moving prey. (B) Animal strikes on the screen are analyzed and sent in real time to the prey application and the reward module (C). (C) An automated dispenser provides rewards (including live rewards) for pre-defined touchscreen strikes. (D) A camera interface and spatial calibration module allows recording and analyzing data from multiple synchronized behavioral cameras. (E) Pose estimation and strike detection models are fed with images from the arena in real time (or offline) and provide feedback to the prey application for enabling predator-prey interactions. (F) A web based user interface allows scheduling automated experiments as well as monitoring and manipulating experiments remotely. (G) All arena components are synchronized internally and with external devices through arduino TTL outputs. (H) Experimental information is logged in and displayed by offline modules for visualizing animal performance. (I) A scheme of the experimental arena. The arena is built with off-the-shelf aluminum profiles. A touch screen is integrated in one of the walls in addition to cameras and a modified automated feeder that are anchored to the top frame.

## Results

### PreyTouch features and performance

Performing prey catch experiments require a repetitive time consuming effort. In order to streamline such experiments and scale up the number of trials, we built a low-cost (Table 1, Methods), automated arena (Figure 1I, see Methods for details on the construction, operation and features of the arena) and wrote a dedicated software suite named PreyTouch. The system’s simple and low-cost design allowed us to easily construct multiple arenas for simultaneous recording of several animals for many consecutive days during which prey catch sessions were periodically initiated. For controlling the session times during these long experiments, we incorporated a dedicated scheduling tool (Methods). Each session was composed of a series of trials with a single prey item (Figure S1B) and a chosen prey dynamics (Figure S1C) presented using the prey application (Figure 1A, Methods). We designed the application to provide animated and visually customizable stimuli mimicking live prey (Figure S1A, S1B). Further, we integrated an interface to PsychoPy to further expand the range of possible visual simulations (Methods). We designed a GUI for configuring all experiment parameters, for example, the quantity or frequency of sessions, the prey types and their movements, the rewards or the light/dark cycles (see Methods, Figure S4F). Finally, we added a system for automatically delivering food rewards (including live prey, (Eisenberg and Shein-Idelson, 2024)) upon successful performance and an alert system for notifying the user in case rewards needed to be replenished or upon system failures.

**Table 1.** List of arena hardware components and prices.

| Component | Description | Price Per Unit (USD) |
| --- | --- | --- |
| Arena frame | Aluminum profile | \$12/meter |
| Arena frame connectors | Nuts, bolts, washers and angle brackets |  |
| Arena walls & floor | 2 X Alucobond sheet (122cm x 244cm gray matte) cut to 5 parts to fit arena walls | \$98 |
| Front & top camera | 2 x Firefly FFY-U3-16S2M-DL | \$598 |
| Front & top camera lens | Boowon 6" - BW60BLF | \$17 |
| Back camera (color) | FLIR Blackfly - BFS-U3-16S2C-CS | \$364 |
| Back camera lens | Computar A4Z2812CS-MPIR, 2.8mm-10mm 1/2.7", CS mount lens | \$109 |
| Camera holder | NOGA Modular holders LC6100 | \$124 |
| Arena computer | Intel Core i7-11700K CPU, 32GB DDR4 memory, NVIDIA GEFORCE RTX 3080 Ti GPU, 500GB SAMSUNG 980 M.2 NVME SSD, and a 2TB 7200 RPM HDD | \$3,245 |
| Arduino Microcontrollers | Arduino Nano Every | \$20 |
| Relay board (lights and heat lamps) | Pololu Basic SPDT Relay Carrier with 5VDC Relay (Assembled) - Omron G5LE-14-DC5 SPDT | \$9 |
| Reward feeder | EVNICE EV200GW fish feeder | \$51 |
| LED Strip | 4 meter, 12V Cold-White approx. 1A/meter | \$20 |
| Lamp Holder | MULTICOMP G6-R32HKY-12 lamp holder | \$3 |
| Temperature sensor | Plastic casing water-proof DS18B20 temperature sensor | \$22 |
| Feeder motor driver | ULN2003 stepper motor driver board | \$4 |
| Led strip PSU | 12V 5A AC-DC power supply | \$13 |
| Feeder PSU | 5V 500mA AC-DC power supply | \$5 |
| Touchscreen | Dell P2418HT; 53x30cm, 24"; Horizontal frequency: 30-83 Hz, Vertical frequency: 50-76 Hz. Resolution: 1920x1080, refresh rate: 60Hz | \$581 |
| sum | | \$5331 |

Measuring strike dynamics (Schaerlaeken et al., 2007) can provide valuable information about the animal’s prey-capture strategies. Additionally, monitoring the predator’s location and posture throughout the experiment (before and after sessions) allows placing prey catch within a wider behavioral context. For this reason, we integrated into PreyTouch modules for video acquisition, calibration and analysis (Methods, Figure 1D). We added a GUI for multi-camera calibration (based on Churuko markers) that allows converting all videos to accurate 2D coordinates within the arena (Figure S2, Methods). In addition, we incorporated a video acquisition GUI allowing camera control (Figure S4D). Finally, we implemented two pose estimation models for extracting movement throughout experiments (Figure 2A): Models for continuous real-time analysis (e.g. for triggering visual stimulation or external devices like heaters or lights) and slower offline models for increasing classification accuracy (Figure 2A, 2B; Methods).

**Figure 2.**
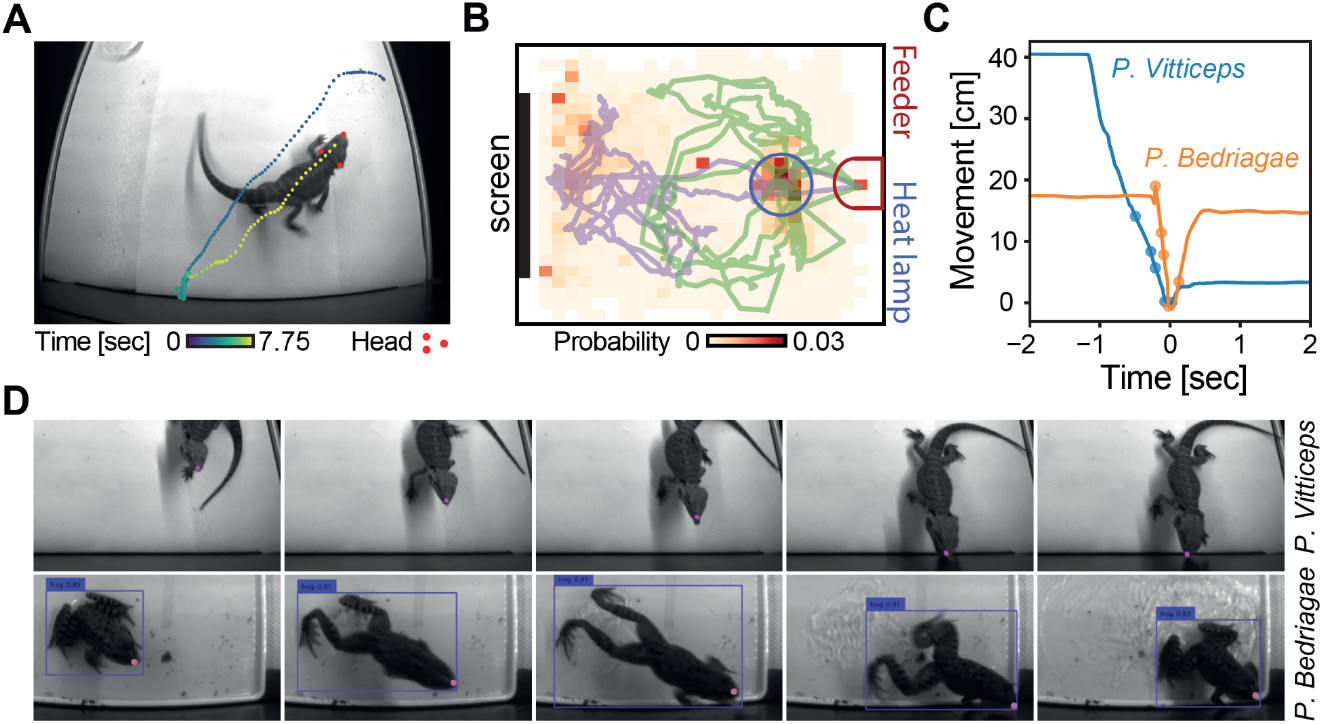
PreyTouch integrates pose estimation for measuring strike dynamics and head and body movement. (A) Top view of a lizard after a strike on its way to receive a reward. Head triangle (marked by red dots) is estimated using DeepLabCut and enables tracking of head movement. Color dots depict the position of the jaw tip as a function of time (color coded). (B) Long-term position tracking of a lizard in the arena. Two trajectories (green and purple) acquired during two 30min epochs of spontaneous behavior. The occupancy probability distribution of the lizard’s position accumulated over 167 hours within one month from one animal is color coded in the background. The feeder location is marked in red and the heat lamp location in blue. (C) Strike dynamics of the animal’s jaw tip (pink dot in (D)) along the axis perpendicular to the screen. The dots correspond to the time of frames in (D). (D) A series of snapshots during prey strike for a lizard (*P. Vitticeps*, top) and a frog (*P. Bedriagae*, bottom). Pink dots depict jaw tips (obtained using a pre-trained DeepLabCut model) and blue rectangles depict a bounding box (obtained in real time using a Yolo5 model).

We used these models to track the animal’s head position and direction (Figure 2A). During the task, this information is useful for tracking the strike dynamics and its relation to prey motion. Figure 2D shows the strike dynamics measured for the lizard *P. Vitticeps* and the frog *P. Bedriagae.* While in both species strikes were characterized by acceleration during approach and deceleration before hitting the target, differences between species were apparent (Figure 2C). Examining head movement trajectories revealed that lizards approached the screen also when sessions were not active and no prey items were displayed (Figure 2B). Integrating these movements over longer times revealed a generally higher occupancy near the screen suggesting that prey catch affected the lizard’s behaviors outside the task (Figure 2B).

Predator-prey interactions are often a bi-directional processes during which both prey and predator modify their behavior in response to the other’s actions. Facilitating such interactions under laboratory experimental settings requires a closed loop system. We therefore incorporated an efficient multiprocessing architecture for executing real-time visual models designed for strike detection. We used this functionality to simulate prey jumps in response to an attack by the predator (Szopa-Comley and Ioannou, 2022). One way to achieve this is to identify the position of the animal (Figure 2A - blue box) and move the prey when the animal is proximal to the screen. However, such proximity does not necessarily indicate a strike attempt. An alternative approach is to identify a distinct pre-strike marker exhibited by the predator. For instance, some predators, like *P. Vitticeps* extend their tongues just before launching an attack (Figure 3A)(Meyers and Herrel, 2005; Schaerlaeken et al., 2007). By training a model (Methods) to discriminate between pre-strike lizard images with extended tongues (Fig. 3A) and images with retracted tongues (Figure 3B), we created a simple yet effective real-time classifier for strike attempts. We achieved a low rate of false positives and false negatives (Figure 3C) as well as AUC (area under curve) close to 1 (Figure S3) when running the classifier on a test set.

**Figure 3:**
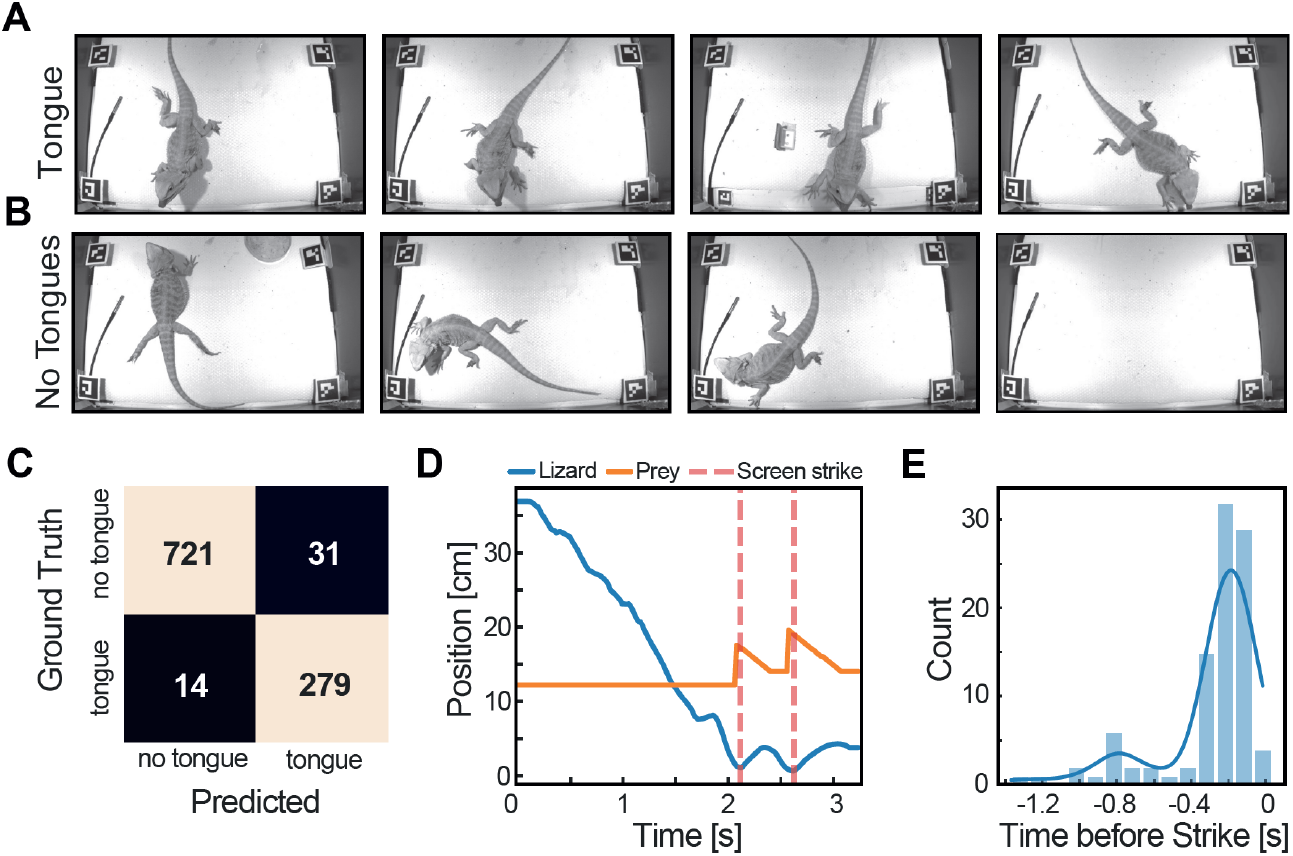
Real-time strike detection and visual feedback for bi-directional predator-prey interactions. (A,B) Sample frames used to train a deep learning model to identify the initiation of lizard strikes. The dataset included two categories: (A) frames capturing a visible tongue just before a strike begins, and (B) frames without visible tongues. (C) Confusion matrix for the model’s performance on the test set showing a very low number of false positives and false negatives. (D) Example of strike detection and visual real time feedback during capture of a horizontally moving prey. Concomitant with the lizard’s approach (blue trace marking jaw tip position on the axis perpendicular to the screen), the detection of tongue protrusions triggers prey jumps (orange curve marking the movement of the prey) occurring just before the lizard strikes the screen (dashed line). (E) A histogram of the strike detection times relative to the screen strike time (t=0). Blue line marks the kernel density estimation for the distribution. The large majority of strike detections occur between 0 - 400 milliseconds prior to the strike.

We used this classifier for a feedback experiment in which a prey moved along a horizontal trajectory on the screen (Figure S1C) and jumped up every time the lizard attempted to strike it. An example of the lizard’s approach towards the screen, followed by two strike attempts is observed in Figure 3D. PreyCatch successfully responded with a prey jump before the lizard hit the screen, as evident from the strike-triggered histogram of prey jump times which occurred ∼200ms before strike time (Figure 3E).

### Prey catch preference and learning in *Pogona Vitticeps*

To demonstrate the performance of PreyTouch we performed a series of prey catch experiments. We focused our analysis on two prey movement patterns: linear horizontal movement and circular movement (Figure 4A). We first examined the engagement of the lizard with the artificial prey stimulus. Following prey onset, lizards sharply moved their head towards the prey (Figure 4C) indicating prey perception. This movement was followed by a quick approach towards the screen (Figure 4C). To further quantify head dynamics relative to the prey, we defined the prey deviation angle (Θ) as the angle between the prey position on the screen and the head direction (Figure 4B; 0° - the head pointing directly at the prey; positive value - left eye on the prey; negative value - right eye on the prey). Following previous reports of gaze lateralization (Frohnwieser et al., 2019; Robins et al., 2005), we compared between sessions in which a prey moved horizontally from right to left (and therefore appeared first with higher probability on the right eye) and vice versa. We found similar movement characteristics in both cases, with symmetric approach patterns (Figure 4C, D). While animals differed in the time it took them to turn their heads towards the prey and approach it (Figure 4E), we didn’t find any evidence for consistent differences in head movement as a function of prey directions which would have indicated visual lateralization. To verify this, we averaged the deviation angles (Θ) during the first second following prey appearance in each trial and tested if they significantly and consistently deviated from 0° to a particular direction (Θ>0° indicating left-eye lateralization and<0° indicating right-eye lateralization). We found that for two out of five tested animals, Θ did not significantly (p<0.05) differ from 0° (1-sample t-test; t=-2.33,p=0.21; t=5.73,p=0.06), and for others, the lateralization patterns were not consistent towards a specific eye (1-sample t-test; t=-56.9,p<0.001; t=19.84,p<0.001; t=11.68,p=0.02).

**Figure 4.**
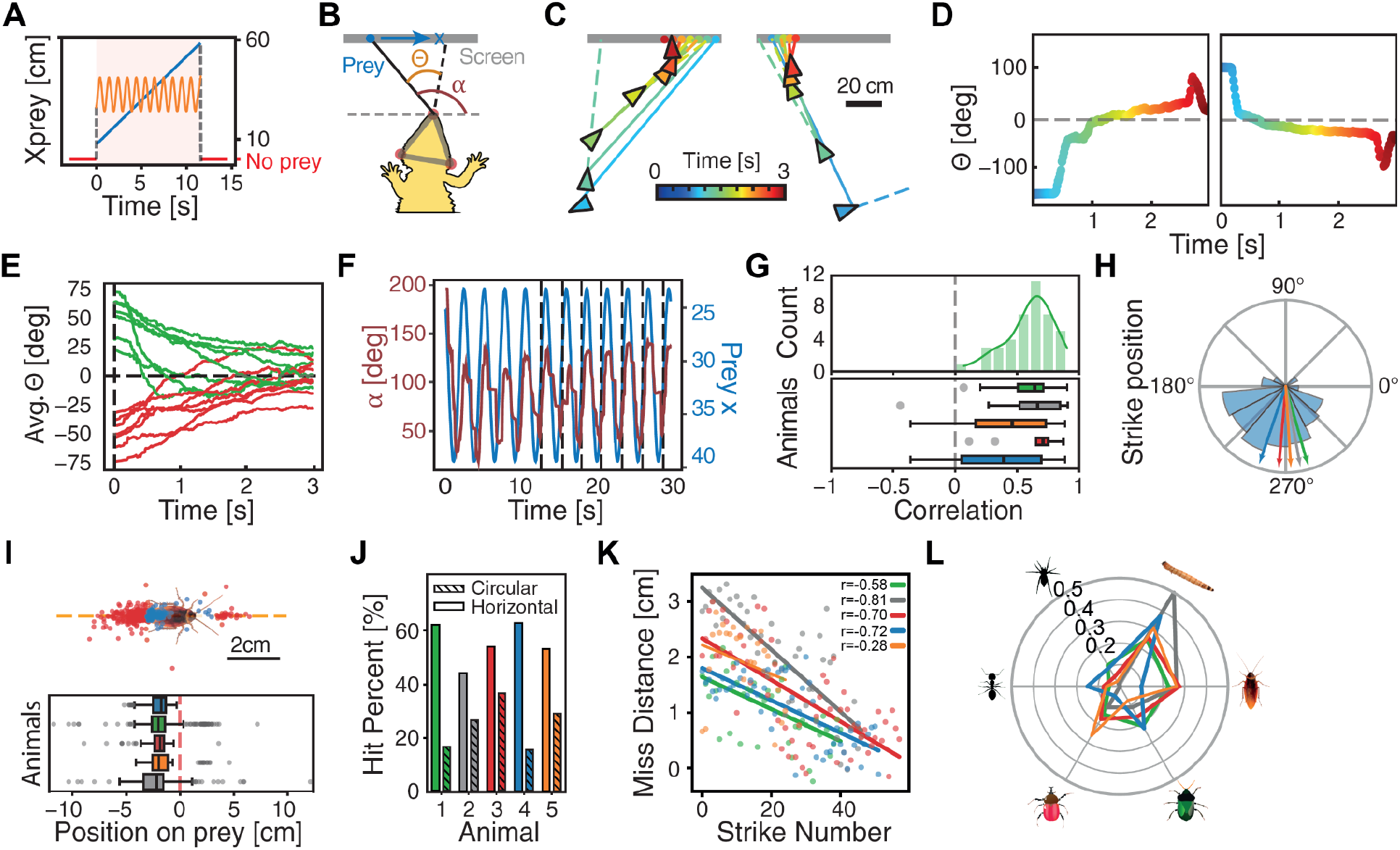
Lizards exhibit prey following, prey learning and prey preferences during prey catch. (A) Dynamics of the prey’s horizontal position on the screen in horizontal (blue line) and circular (orange line) movement types. (B) A schematic illustration of a lizard during prey catch. Head position is marked by a triangle (shaded gray) whose direction (⍺) is determined by three head points (red dots, see also Figure 2A). The prey is marked by a blue dot. Θ (prey deviation angle) marks the angle between the head direction (black dotted line) and prey direction (black solid line). (C) Head and prey dynamics. Head position (triangles), prey position (filled dots), head direction (dashed color lines) and head-to-prey direction (solid lines) for single frames (every 20^th^ frame is shown) color coded for time (t=0 prey appearance) during a single trial of prey catch. Two trials with prey movement to the left (left) and to the right (right) are shown. Note change in head direction following the appearance of the prey on the screen and the consequent movement towards it. (D) The deviation angle (Θ) dynamics for the trials in (C). Same color code as in (C). (E) Average deviation angle (Θ) dynamics after prey appearance (t=0) for trials with prey moving horizontally from right to left (red) and from left to right (green). Lizards (n=5) converge towards binocular gaze on the prey (Θ=∼0°) within a few seconds of prey onset. (F) Head angle, ⍺ (red line), and prey’s horizontal position (blue line) as a function of time in circular trials. Black vertical-dotted lines indicate strike attempts. Notice locking of the head to prey motion. (G) Distributions of correlation values between the head angle and the bug’s horizontal position (as in F) during circularly moving prey trials in one animal (top) and all animals (bottom; n=5). Box plots indicating the 25th-75th percentile (middle line marks the median). Whiskers indicate the 0th and 100th percentile and outliers are marked as gray dots. (H) Distribution of prey position (angles on the circle) at strike times during circular movement sessions for one animal (blue). The average strike angle in different animals is marked by arrows. (I-Top) Scatter plot of screen strikes recorded from a single lizard (cockroach prey). Misses (miss distance > bug body radius, 1.85cm; n=316; red) and successful rewarded hits (n=129; blue) are shown. (I-Bottom) Strike hit statistics during circular movement trials for 5 lizards (box plots as in (G)). (J) Percentage of successful strikes for horizontal and circular movement trials across animals. Notice higher success rates for horizontal movement. (K) Dynamics of miss distance for consecutive strikes in circular trials for all animals showing gradual improvement quantified by significant negative correlation (legend of the left: r, correlation coefficient; p, p-value; r=-0.58,p<0.001; r=-0.81,p<0.001; r=-0.7,p<0.001; r=-0.72,p<0.001; r=-0.28,p=0.182). (L) Polar plot of normalized (across prey types) prey choice probabilities (fraction of strikes per number of presented trials). Plots (G)-(L) show statistics from 5 lizards with consistent color coded identity.

We next examined head dynamics during circular movement. In these trials, the prey stayed on the screen for longer periods. We observed that the lizards moved their head (quantified by the head angle, ⍺, Figure 4B) to follow the prey for several cycles before striking (Figure 4F). This movement resulted in positive correlation values between the prey position and head movement (Figure 4G - top). Such correlations were consistent across animals (Figure 4G - bottom). Interestingly, although lizards observed and tracked the prey during all phases of the bug’s circular motion, they only stroked the prey when it reached the bottom of the screen (prey rotation angle of 270°±60°, Figure 4H).

To study the lizard’s prey catch accuracy, we analyzed the positions of strikes relative to the prey. Strike positions were registered on the screen in real time and successful strikes (defined by falling within a predefined radius surrounding the bug center; blue dots in Figure 4I - top) were rewarded with a live mealworm delivered by the automated feeder. The strike position was asymmetric with respect to the prey direction of movement with most strikes located in the rear part of the prey (Figure 4I - top). This strike pattern was consistent across lizards (Figure 4I bottom). Lizards managed to successfully strike the prey in a large fraction of the strikes but success rates were not homogenous across prey movement types (Figure 4J). Lizards were more successful in striking horizontally moving prey and less successful in striking circularly moving prey (Figure 4J) indicating a difference in task difficulty.

To understand if strike performance could be improved, we examined the miss distance (Euclidean distance between the strike position and the bug position) for each registered strike and tracked it over time. We found that lizards significantly decreased their missing distance as virtual-hunting progressed (Figure 4K). The rate of decrease differed across animals, but all showed a similar trend of improvement in catching the circularly moving prey (Figure 4K). Surprisingly, we did not find a significant improvement in the miss distance during horizontal movement trials across animals (n=5 animals; r=correlation coefficient; r=-0.16,p=0.16; r=-0.33,p=0.026; r=-0.14,p=0.195; r=-0.18, p=0.094; r=-0.06;p=0.591).

Finally, we examined if lizards had preference to specific prey items. To do so, we presented prey items differing in shape and color. We found that different prey items elicited a different number of strike attempts. The preferred prey item (with the highest fraction of strike attempts per trial) was mealworms and the second preferred item was a cockroach (Figure 4G). Interestingly, the main live food provided to the lizards both before the experiments (in their home cages in the animal house) and as reward during the experiment was mealworms (*T. Molitor* or small *Z. Morio*). The lizards occasionally received cockroaches (*N. Cinerea*) as live feed in the animal house and never received any of the other prey items.

## Discussion

In this manuscript we introduce a novel experimental toolkit for conducting prey capture experiments using a touch screen. PreyTouch is constructed from off-the-shelf low-cost components increasing its accessibility to a variety of laboratories. The open-source nature of PreyTouch and its integration with PsychoPy empowers experimenters with easy customization for adapting the system according to specific requirements and preferred species. These properties together with its real-time processing, the long experiments scheduler and its compatibility with electrophysiological recordings make PreyTouch a valuable tool for comparative studies across species.

PreyTouch is suited for studying animal models capable of catching prey by touching a screen. This approach diverges from previous research utilizing live prey (Borghuis and Leonardo, 2015; Michaiel et al., 2020; Schaerlaeken et al., 2007). The main advantage of using live prey is its realistic appearance which more likely elicits natural predator-prey interactions. However, this advantage comes at the price of limited prey control. For example, trial to trial variability in live prey movements may change the animal’s capture strategies thus limiting reproducibility. Further, it is difficult to accurately manipulate a live prey’s appearance, position and dynamics. This ability to pre-determine prey parameters is important for using prey catch paradigms to systematically study sensory processing and movement-dependent neural computations (Yoo et al., 2020). Finally, when using live prey, the number of prey catch trials is limited by the satiation of the animal. An alternative to live prey is the usage of robots mimicking prey items (Mischiati et al., 2015; Szopa-Comley and Ioannou, 2022). However, such robots require substantial design efforts, complex arenas and are hard to generalize to different species.

In contrast to using live prey or robots, PreyTouch allows full control over prey properties and easy placement of prey backgrounds and visual context (Figure 1, S1), thus offering a good tradeoff between realism and control. In addition, the low latency real-time feedback integrated in PreyTouch exemplifies how prey-predator interactions can be integrated in experimental designs (Figure 3). Finally, the integration of the animal’s pose (extracted from recorded video) with the touchscreen strike logs offers an accurate quantification of the fine structure of animal movements and choices (Szabo et al., 2021) (Figure 2,4). To the best of our knowledge, no other system offers these features for prey catch experiments. Importantly, the above benefits in prey control are valuable as long as animals are motivated to try and catch virtual prey on the screen. Our experiments revealed that lizards learn to associate between striking a prey on the screen and the delivery of a real prey on the opposite side of the arena (Figure 4). This may have been the reason for the consistently robust prey catch behavior that allowed us to collect many trials (Figure 4).

The proof of concept experiments we performed suggest that PreyTouch can be used as a platform for studying learning, and provide insights on how reptiles (and other animals) adjust innate behaviors to accommodate changing environments (Figure 4C, 4F, 4K). For many years, reptiles were considered to be slow, adamant, and with limited learning capacity due to their underperformance in cognitive tasks (De Meester and Baeckens, 2021). Using PreyTouch, we observed significant and fast improvement in the performance of *P. Vitticeps* lizards during circular prey-catching trials (Figure 4K). This result is consistent with the idea that achieving fast learning in reptiles strongly depends on task design (Burghardt, 1977, 2013) highlighting the advantages of the prey catch paradigm. In contrast to the improvement with the circular prey, no learning was observed in horizontal movement trials. This may be due to the lower difficulty of this task, as supported by the higher success rates we measured for catching horizontally moving prey (Figure 4J). This lower difficulty may have decreased the lizard’s motivation to change strategy and improve (Ashwood et al., 2022). We also observed that lizards discriminated between different types of prey (in line with reports of complex visual processing abilities (Qi et al., 2018; Szabo et al., 2018; Szabo and Whiting, 2022)), and preferred those found in their regular diet (Figure 4L). However, lizards also performed strikes on novel prey that they did not encounter before (Figure 4L) suggesting lizards can balance between novel and familiar prey choices. Such a property can be important if diversifying prey appearance is required in the experimental design.

PreyTouch provides opportunities for performing long-term automated experiments. Behavioral experiments with reptiles tend to last longer and require more repetitions, possibly due to their slower metabolism which diminishes the effectiveness of food as a motivator, lower curiosity or diminished play behavior relative to mammals and birds (Burghardt, 1977, 2013; Wilkinson and Huber, 2012). Following previous success with automated approaches (Eisenberg and Shein-Idelson, 2024), we addressed these caveats by integrating the PreyTouch scheduler agent allowing automatic and remote monitoring of animal progress as well as optimizing experiment schedules. This strategy enabled the collection of large prey catch datasets with minimal effort while animals remain engaged for prolonged periods of time (Figure 4). Further, the low-cost (Table 1) and simple design allows replicating the experimental setup thus further increasing experimental throughput.

Moving forward, PreyTouch offers opportunities for studying new avenues in animal cognition. For example, the system could be used to explore the visual processing capacity or the learning capacity of animals. We can ask to which degree can lizards link specific prey items or specific movement types with rewards. We observed that while lizards continuously track prey movement (Figure 4F, 4G), they strike mostly at specific strategic points (lower part of the screen), possibly to maximize their success (Figure 4H). Such a strategy hints that lizards possess an “understanding” of their prey’s dynamics and may thus possess predictive behaviors (McNamee and Wolpert, 2019). Such behaviors can be examined by manipulating prey movement statistics and measuring strike success dynamics to understand if such statistics can be learned. For example, one could examine if lizards use information about prey velocity or prey spatial distribution and modify their strike statistics accordingly (Borghuis and Leonardo, 2015). Using PreyTouch, such questions could be investigated across animal classes (Figure 2A). The combination of flexible manipulation for addressing various questions with the ubiquity of prey catching behavior suggests PreyTouch as a valuable platform for comparative studies of animal cognition across species which may in turn provide valuable insights on brain evolution (Brenowitz and Zakon, 2015; Laurent, 2020; Yartsev, 2017).

## Materials and Methods

Below we provide details about the construction and features of the PreyTouch system. All code and arena construction instructions can be accessed on GitHub at the following URL: https://github.com/EvolutionaryNeuralCodingLab/PreyTouch

### Hardware

The experimental arena was constructed from off-the-shelf low-cost components (Figure 1I, Table 1). These components include an arena cage, a touch screen, cameras, a server, and arena peripherals and are described in detail below.

#### Arena Cage

The arena is built from a frame of connected aluminum profiles with dimensions of 70×100×45cm (WxLxH) (Figure 1I). Arena walls and floor were made from 3 mm thick aluminum composite panels (Eisenberg and Shein-Idelson, 2024). The choice of conductive wall is optimized for experiments with electrophysiological measurement requiring a faraday cage.

#### Touch Screen

A touch screen (Dell P2418HT; 53×30cm, 24”; Horizontal frequency: 30-83 Hz, Vertical frequency: 50-76 Hz. Resolution: 1920×1080, refresh rate: 60Hz.) was integrated into one of the arena walls, and cameras as well as a modified and automated worm feeder were anchored to the arena top rail (Figure 1I). For experiments combining electrophysiology we used a low noise touch screen (ELO Touch Solutions AccuTouch 1790L 17” LCD open frame) that did not induce 50Hz artifacts when animals got close to the screen.

#### Cameras

Two sets of cameras were used with PreyTouch and are fully supported by the system: (1) FLIR - Firefly FFY-U3-16S2M-DL + Blackfly S BFS-U3-16S2C, (2) Allied-Vision - 1800 U-158c + 1800 U-158m. Cameras were anchored to the arena’s top aluminum profiles using adjustable arms (Noga Engineering & Technology, LC6100) and a custom 3D printed connector. Video was recorded at a frame rate of 60hz.

#### Server

PreyTouch was installed on a linux desktop computer (ubuntu 22.04) with the following specs: CPU: Intel Core i7-11700K, memory: 32GB DDR4, GPU Nvidia GeForce RTX 3080Ti (nvidia-driver: 530.30.02, cuda: 11.7), storage disks: 500GB SAMSUNG 980 M.2 NVME SSD + 2TB 7200 RPM HDD.

#### Arena peripherals

All peripheral devices, including the reward dispenser, LED lights, temperature sensors, and camera triggers, are interfaced using an Arduino (Nano Every). The complete code for both the Arduino and the service facilitating communication through MQTT were sourced from the ReptiLearn project **(Eisenberg and Shein-Idelson, 2024)**.

#### Lights

The arena was lit using an LED strip (12V, 6500K white LEDs, approximately 3.4 meters long) that was attached using adhesive to profiles at the top edge of the four arena walls. This provided relatively uniform lighting across the arena, minimizing shadows. The strip was controlled using a relay module (based on an Omron G5LE-14-DC5 5VDC SPDT relay). The module’s EN, VDD, and GND control ports were connected to one of the Arduino boards that sent on/off TTLs to control the DC output connected to the LED strip’s power supply unit (12V, 5A).

#### Live prey dispenser

For rewarding animals with live prey we utilized a readily available aquarium feeder (EVNICE EV200GW), which was attached to the arena’s structure using its built-in clamp. The feeder was modified to decrease its response time and driven by an Arduino board **(Eisenberg and Shein-Idelson, 2024)**. Feeders were stocked with worms, each housed in individual compartments (15 in total) with sufficient food for gut-loading. To ensure the animals remained eager for the rewards, smaller worms proved more effective, leading us to choose *Tenebrio molitor* larvae. In instances requiring a larger worm supply between refills, we optimized space by vertically arranging the feeders, achieving capacities of up to 45 worms in three vertically aligned feeders. Feeders stacked vertically were positioned such that the upper feeder dispensed rewards through the release aperture of the feeder below it. The experimental module kept track of the reward inventory in each feeder to determine which unit should release a reward next.

#### Temperature sensors

We employed two DS18B20 digital temperature sensors, each encased in plastic, to monitor the temperature within the arena. The first sensor was affixed to the arena’s rear wall to gauge ambient temperatures, while the second was positioned at the base of the wall opposite, directly beneath the heat lamp in the basking area. These sensors were linked to an Arduino board for data collection and 5V supply.

### Software

#### PreyTouch Software

The majority of the PreyTouch codebase was developed and tested using Python 3.8. To enable concurrent processing with multiple cameras, the system employs a multiprocessing architecture with a Redis store. The software leverages Flask API (v2.2.2) for execution and hosts a user-friendly management web interface (Figure S4) created using jQuery (v1.11.0). PreyTouch is compatible with touch screens and has the capability to exert full control over them, facilitating the display of the prey application (as detailed in the subsequent section), and monitoring and logging screen touches. Communication within the system, such as interactions between camera processes, the API, the prey application, and peripheral components, is established through the MQTT messaging protocol. For instance, commands are transmitted to operate the feeder dispenser (reward) or to receive instructions from the server (e.g., initiation of bug trials or other events like bug jumps). Via the MQTT protocol, the application systematically communicates a comprehensive log of all behavioral events, encompassing screen touches and bug trajectories.

#### Prey Application

A prey application (Figure 1A) was designed to provide realistic visual stimuli mimicking live prey. To achieve this, each prey item was constructed of a sequence of still images that were repeatedly displayed to create animation (Figure S1A) in addition to an image displayed following a successful hit (Figure S1A - left). Prey items can be selected from an existing repertoire (Figure S1B) or provided as input for seamless customization of new prey items. In addition to the appearance of prey, its dynamics is also configurable. Movement profiles can be selected from a range of implemented profiles (Figure S1C) or customized by providing the (x,y) trajectory. In addition, the application allows determining parameters such as movement duration, velocity or prey size. Finally, static objects (e.g., walls, obstacles) can be added if necessary (Figure S1C). The prey application receives input from the touch screen and responds to animal strikes accordingly (Figure S1A). The Prey application is written in Vue.js, a JavaScript framework that is well-suited for web-based (browser independent) animation with asynchronous events required for fast responsiveness. In addition, the application is based on MQTT messaging, allowing it to efficiently communicate with devices. This design scheme allows fast updating of prey appearance following a successful strike, sending commands to operate the reward dispenser or receiving commands from the server. The MQTT protocol is also used to send a full log of all behavioral events, such as screen touches and prey trajectories.

#### Logging and Databases

The system performs logging (Figure 1H) through two mechanisms: storing data in local files and a designated database. PreyTouch utilizes PostgreSQL as its chosen database, with communication facilitated through SQLAlchemy (v1.4.39). Additionally, the system incorporates a migration method to the database using Alembic (v1.8.1). Various data types are consistently saved to both the database and local files: (1) videos from the arena (actual videos are saved locally, while frame times are saved to both local files and DB), (2) screen touches, (3) bug trajectories on the screen, (4) experiment timings and configuration, (5) temperature in the arena, (6) off-line analysis, (7) rewards. For more information on the structure of the database please refer to Figure S5.

#### Detection Models

Monitoring predator dynamics throughout the experiment is important for gaining a comprehensive understanding of prey catch behavior. For this reason, we integrated into PreyTouch modules for automated video acquisition from multiple cameras (Figure 1D), in addition to pose estimation models for extracting movement information (e.g. DeepLabCut **(Mathis et al., 2018)**). These models can operate in two modes: (1) In real-time mode, video frames are continuously analyzed for activity based triggering of visual stimulation or external devices like heaters or lights. (2) In offline mode, deeper and slower models are utilized (when the system is not actively running experiments) for increasing classification accuracy. Support is already integrated for a range of models, including DeepLabCut, YOLO, and any models based on the PyTorch framework.

#### Tongue Model - Real-Time strike detection

The tongue detection model employs ResNet18 as the backbone to embed input images from 224×224×3 parameters to 512 parameters. The model, developed in PyTorch (v2.0.1), was designed as follows:

(resnet): resnet18(in_features=150528, out_features=512)

(fc1): Linear(in_features=512, out_features=120, bias=True)

(fc2): Linear(in_features=120, out_features=60, bias=True)

(fc3): Linear(in_features=60, out_features=2, bias=True)

(dropout): Dropout(p=0.2, inplace=False)

(norm): BatchNorm1d(512, eps=1e-05, momentum=0.1, affine=True, track_running_stats=True)

#### Scheduler and Agent

In order to streamline experiments and scale up the number of trials, we incorporated a scheduling tool in PreyTouch. This tool allows performing automated experiments without user intervention. Before the beginning of the experiment, the arena is configured and parameters such as the quantity, frequency, types of prey-catching experiments, rewards and the light/dark cycles, are set. After configuration, animals can be placed in the system for extended periods of days, thus, allowing the system to accumulate valuable prey catch data. The only required human intervention is filling up the reward dispensers. Further, the system tracks the feeders’ status and sends an alert to the experimenter in case the rewards run out. Alerts are also automatically sent in case of system failures. The scheduler, implemented as a Python module, initializes with the system, conducts checks, and executes scheduled tasks at their designated times. Examples of checks performed by the scheduler include monitoring the state of arena lights, initiating or concluding camera acquisition, commencing scheduled experiments, executing offline analysis, managing database migrations, and compressing videos. Configuration adjustments for the scheduler can be executed either through its module (scheduler.py) or via the configuration file. For detailed information on scheduler configuration, please consult the GitHub link. On the other hand, the agent, guided by a plan outlined in agent_config.yaml, autonomously launches experiments. It actively monitors the animal’s state and performance, adjusting experiments accordingly. For guidance on configuring the agent, refer to the GitHub link.

#### Camera Calibration and Real-World Coordinates

PreyTouch integrates a toolbox for easy camera undistortion (based on checkerboard detection) and for converting pose estimation results to 2D coordinates within the arena (Using ChArUco markers). This tool is based on the openCV package for computer vision (Bradski 2000) and its integrated camera calibration functions. The first step of the calibration process is undistortion and it requires a printed checkerboard (size: 9×6 square corners) that is captured by the camera in various orientations within the arena. The board is detected using the openCV *findChessboardCorners* function. The detected corners of the chess board are then fed to the *calibrateCamera* function which provides the undistort-homography-matrix (see Figure S2 for the result of applying the undistort-homography-matrix on a frame). The resulting undistort-homography matrix is saved by date and camera name, facilitating subsequent offline analyses of frame data. PreyTouch offers a method for extracting real-world coordinates by converting frame detections from pixels to centimeters within a predefined coordinate system in the arena. To achieve this, users are instructed to print a ChArUco board matching the arena’s dimensions and place it within. The detection of the ChArUco board is made by the openCV *cv2.aruco.detectMarkers* function that provides the corners of the detected markers with their marker ID. These detected points are first undistorted using the undistort-homography matrix. Since the board matches the arena dimensions it is possible to calculate the real-world locations of each marker. Using the undistorted locations of the markers and their real-world locations PreyTouch calculates the real-world-projection-matrix using the openCV *findHomography* function, and like the undistort matrix it is saved by camera and date. This procedure is conducted on a per-camera basis, provided the camera remains stationary. In the event of any camera movement, it becomes imperative to repeat this process. For more details on the process and explanations for how to run it in PreyTouch see <u>camera calibration guide</u>.

#### Integration with OpenEphys

PreyTouch was tested with the OpenEphys electrophysiology recording system. The cameras in PreyTouch are set in trigger mode, which is controlled by the PreyTouch software through a camera trigger Arduino. To ensure synchronization with OpenEphys, PreyTouch pauses the camera trigger for 8 seconds at the start and end of each experiment block. This pause allows one to identify these trigger interruptions in the recorded OpenEphys digital data and accurately extract the correct experiment triggers. Additionally, PreyTouch activates the IR LED for 1 second at the beginning of a block immediately after the trigger pause and at the end just before the trigger pause. This IR signal can be used to confirm that the synchronization is functioning correctly by comparing video signals with trigger times.

#### Integration with PsychoPy

Experiments in the visual and auditory domains developed using the PsychoPy framework **(Peirce et al., 2019)** can be easily implemented in PreyTouch. This is achieved by transferring the PsychoPy experiment files to a specific directory within PreyTouch (detailed guidelines are provided on the PreyTouch <u>GitHub repository</u>).

### Prey catch task

Lizards were introduced into the experimental arena that included a heat lamp, a small shelter facing the screen, and a water dish. The experiments began after a 12-hour acclimation period in their new environment. The primary source of nourishment for the lizards came from the feeder dispensers (*Tenebrio molitor* larvae) awarded upon successful virtual insect captures (one worm for each successful attempt). Additionally, the lizards received greens once a week. The experimental activities occurred daily from 08:00 to 17:00, with a series of trials conducted hourly. The experiment involved various trial types, each with specific goals that needed to be met before moving on to the next phase. The system automatically calculated these goals and scheduled the sessions.

Experimental Phases:

1. Random Low Horizontal: Insects move horizontally between holes at random speeds (2, 4, 6, 8 cm/sec) and directions. The objective was to achieve 30 hits at each speed.
2. Circular Movement: Insects move in a circular path with random speeds (2, 4, 6, 8 cm/sec) and directions. The aim was to secure 20 hits at each speed, with a 20% chance of receiving a reward on a miss.
3. Low Horizontal: Insects move at a constant speed of 6 cm/sec horizontally, starting from right to left for 100 trials before switching directions for another 100 trials.

### Analysis

#### Alignment of bugs data and pose

The synchronization of camera clocks with the server clock is lacking, resulting in a misalignment between the extracted animal pose from the camera frames and the screen events obtained from the bug application. To address this disparity, when commencing an experiment, we initiate a query to each camera to retrieve its current time. Subsequently, we calculate the time difference in nanoseconds between the server time and the camera time, preserving this value for each camera process. This calculated difference serves the purpose of converting frame timestamps to align with the server time.

#### Regression models

All regression models in the paper (Figure 4) are calculated using the Scipy (v1.10) method of stats.lingress with 2-sided alternative hypothesis. The lingress method outputs the slope and intercept of the regression model, the Pearson correlation coefficient (r-value) and the p-value for a hypothesis test whose null hypothesis is that the slope is zero, using Wald Test with t-distribution of the test statistic.

#### Evaluation of classification models

All the assessment of classification models, such as tongue-out detection, is conducted utilizing the scikit-learn package (v1.1.2) for metrics. Upon the completion of model training on the train-set, the test-set (observations unseen during training) is subjected to scikit-learn methods to compute the following metrics: Confusion Matrix: A pivotal table utilized in the evaluation of classification models, presenting counts of true positive (TP), true negative (TN), false positive (FP), and false negative (FN) predictions, summarizing model performance comprehensively.

ROC Curve: (Receiver operating characteristic) Offering insights into the sensitivity (true positive rate) and specificity (false positive rate) trade-off across diverse decision thresholds, the ROC curve is a valuable tool for understanding model performance. Precision-Recall Curve: Particularly essential in imbalanced scenarios, the precision-recall curve delineates the trade-off between precision and recall. It aids in threshold selection and provides a comprehensive view of a model’s efficacy in capturing positive instances. The area under the PR curve (AUC-PR or AP) condenses the information from the entire PR graph into a single metric, with a higher AP indicating superior overall model performance, serving as a useful summary measure.

### Animals

#### Pogona vitticeps

Lizards were purchased from local dealers and housed in an animal house at I. Meier Segals Garden for Zoological Research at Tel Aviv University. Lizards were kept in a 12–12 h light (07:00–19:00) and dark cycle and a room temperature of 24 °C. All experiments were approved by Tel Aviv University ethical committee (Approval number: 04-21-055). Prior to entering the arena, lizards (Table 2) are fed a diet comprising mealworms (*Tenebrio molitor*), cockroaches (*Nauphoeta cinerea*), and vegetables such as carrots and parsley. Once introduced to the arena, the lizards exclusively receive mealworms from the automated reward dispenser with a petri dish containing vegetables provided at the end of the week. A water bowl is always accessible.

**Table 2.**
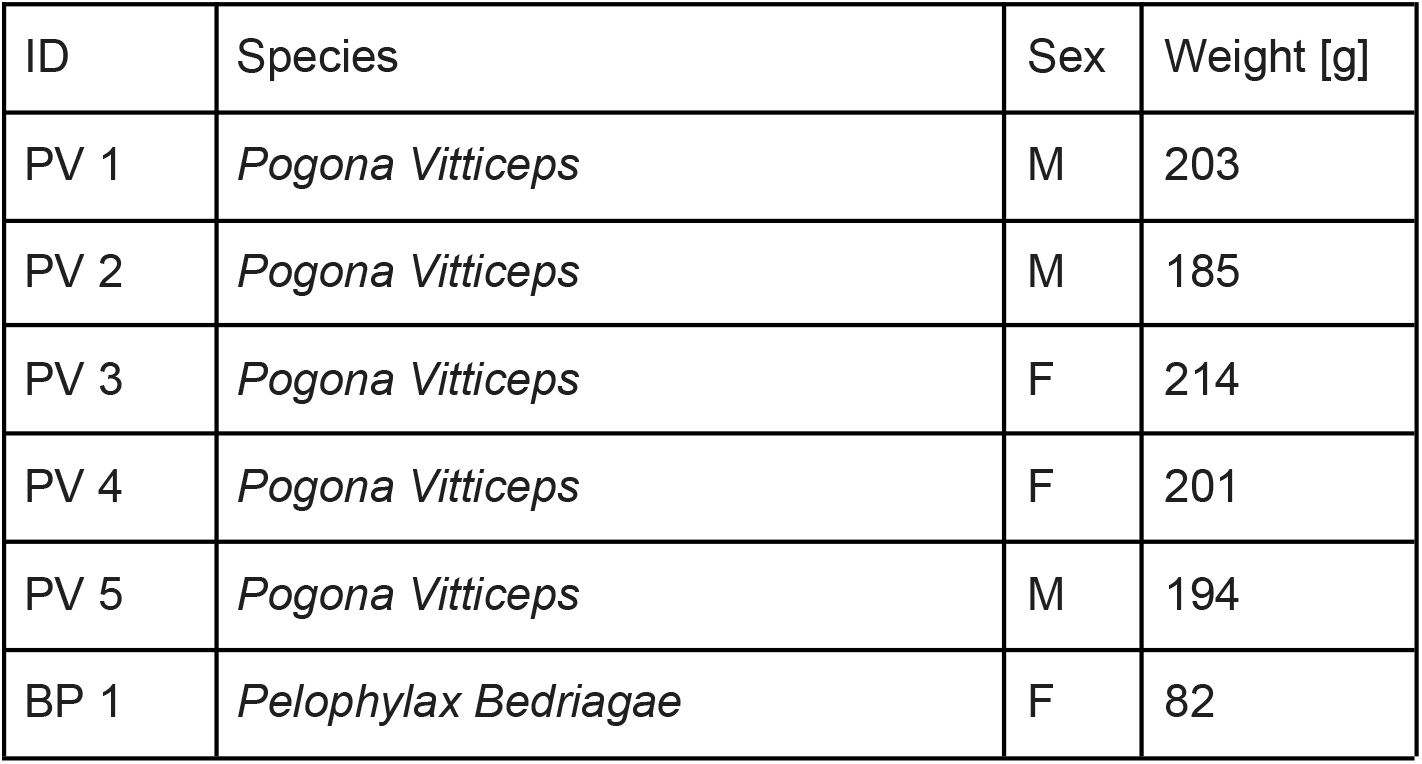
List of animals participating in the study.

#### Pelophylax bedriagae

Frogs were caught from the wild and housed in an outdoor pond at I. Meier Segals Garden for Zoological Research at Tel Aviv University. All experiments were approved by Tel Aviv University ethical committee (Approval number: 2208-155-4). Before introducing the frog to the arena, the top of the arena is covered with a net (preventing the frog from leaping out), and a large plate with dechlorinated water is placed inside the arena in proximity to the screen. Similar to the lizards, the frog was rewarded with mealworms (*Tenebrio molitor.* Also their only type of food outside the experimental arena) from the dispenser upon successful strikes.

## Supplementary Materials

**Figure S1:**
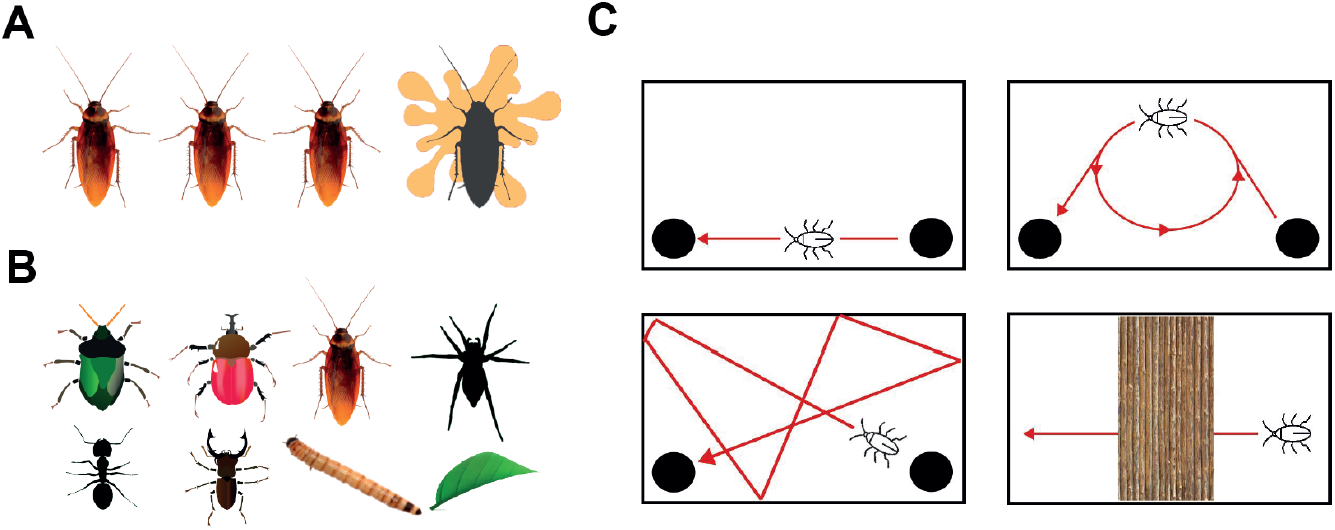
The prey catch application allows easy configuration of prey shape and trajectory. (A) Example of a sequence of input movement images defining prey dynamics. The right image is displayed following successful prey capture. (B) Examples of prey items integrated in the application. (C) Schematics of movement profiles supported by the application. From left to right and top to bottom: linear movement in random directions, circular movement, linear horizontal movement, linear movement under an occluded object. Black lines mark screen boundaries, and black dots are entry and exit holes for the prey. The application also allows customizing prey shapes and movements.

**Figure S2:**
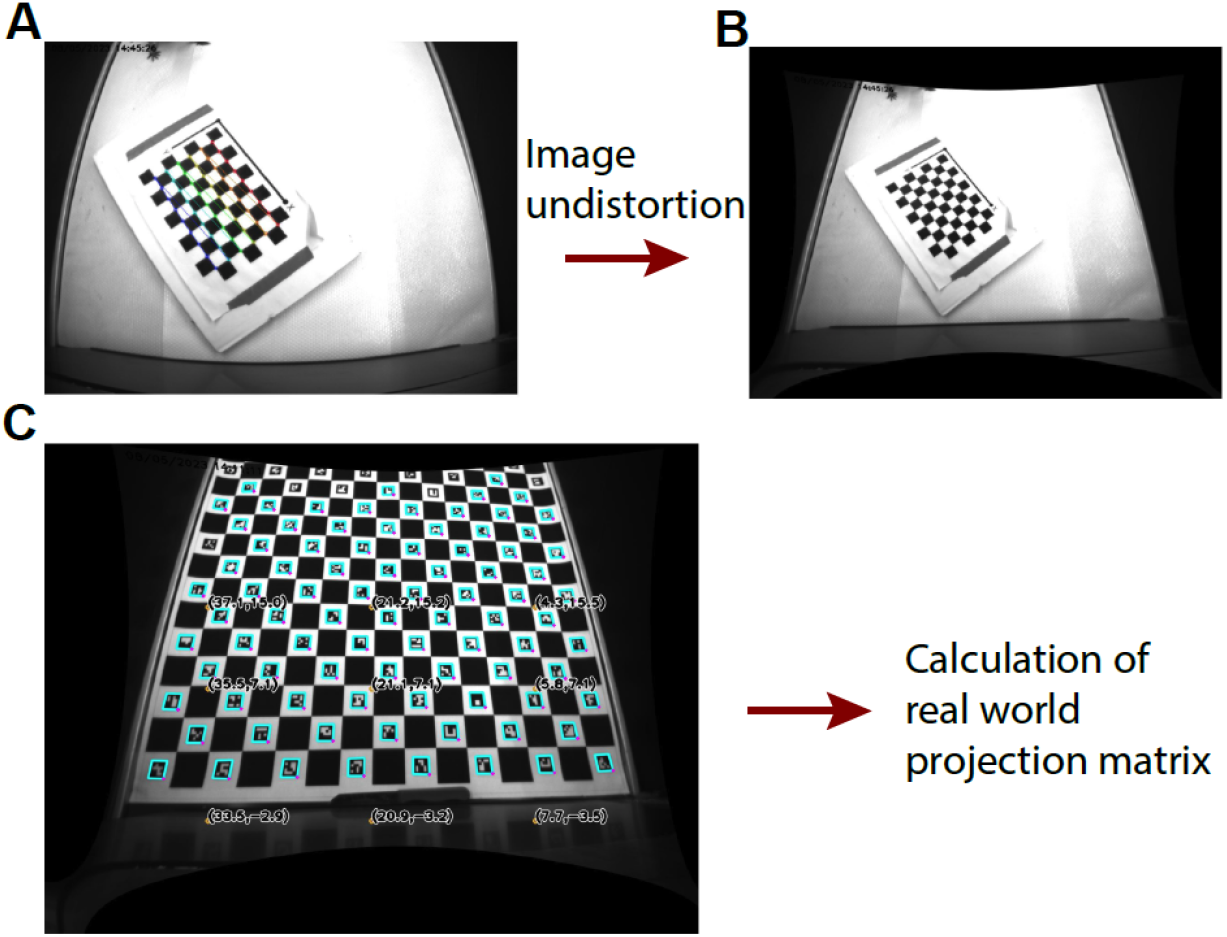
Camera Calibration. (A) Detection of Chess-board by openCV. By detecting several such frames in different orientations of the chess-board, one can calculate the homography matrix to undistort the original frames (B). (C) Detection of Charuco markers on the arena floor (cyan rectangles). The detected corners of the markers are first undistorted using the homography matrix and then are used for calculating the real-world projection matrix that translates the frame coordinates (pixels) to real-world coordinates (cm). 9 examples of projections are written on the frame with their resulting real-world coordinates.

**Figure S3.**
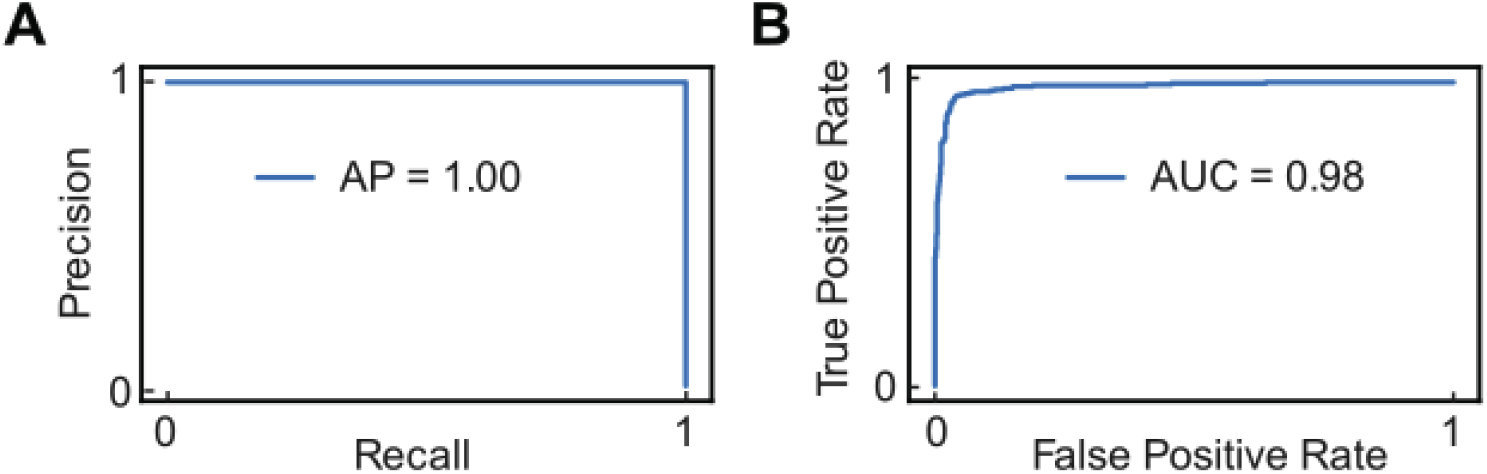
Strike detection real-time model. (A) Precision-Recall graph is essential for understanding the performance of classification models, particularly in imbalanced scenarios. It provides insights into the trade-off between precision and recall, aids in threshold selection, and offers a comprehensive view of a model’s behavior in capturing positive instances. The area under the PR curve (AUC-PR or AP) condenses the information from the entire PR graph into a single metric. A higher AP generally indicates better overall model performance, making it a useful summary measure.(B) ROC curve provides valuable insights into the trade-off between a model’s sensitivity (true positive rate) and specificity (False positive rate) across different decision thresholds.

**Figure S4:**
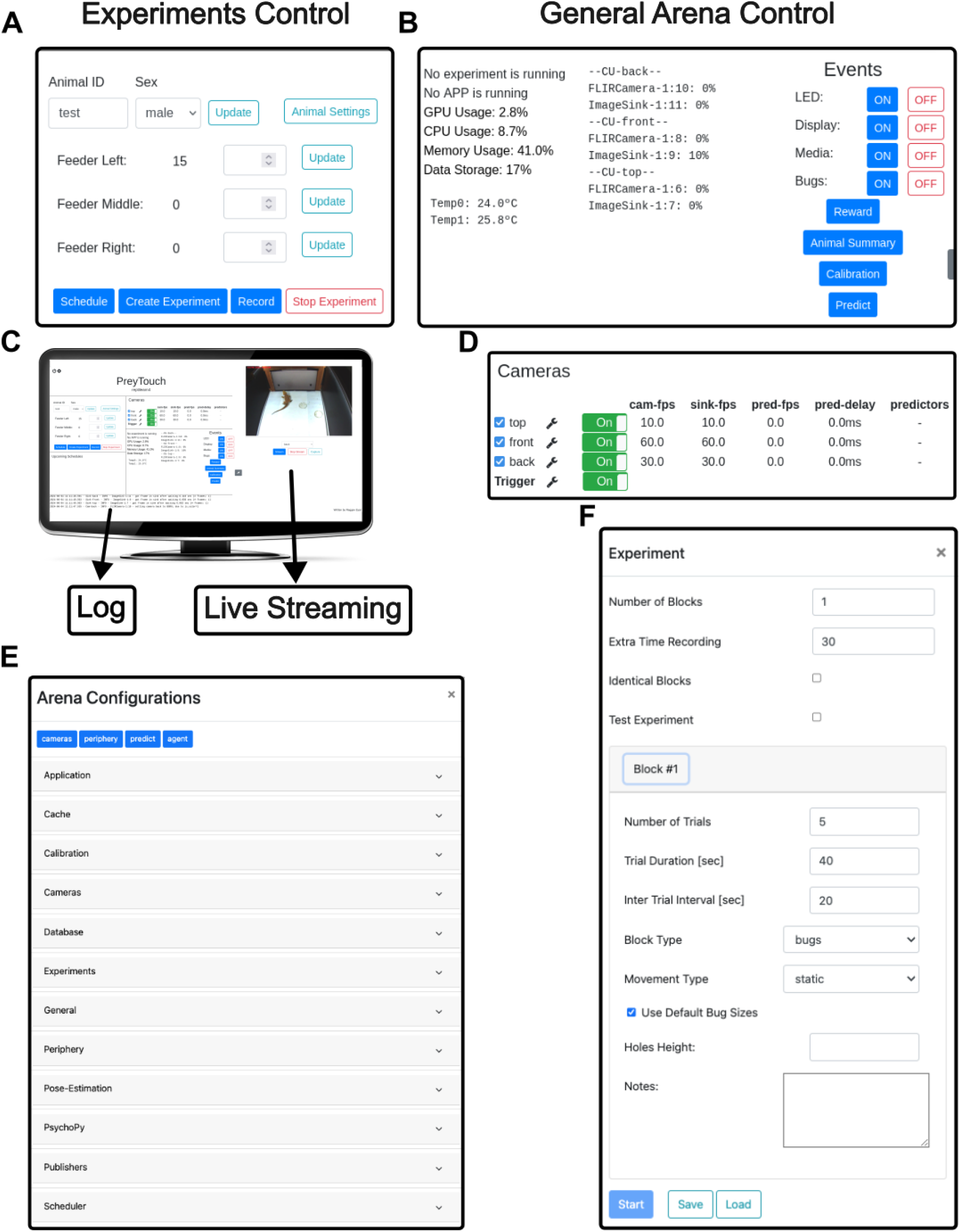
Management WebUI. (A) Experiment panel (left panel; C). Animal information, Feeders loads and shortcuts to scheduling experiment, creating experiments (F) and recordings. (B) General arena control (lower middle panel; C): server metrics, periphery controls and shortcuts to general forms like calibration and predictions. (C) Demonstration of the management UI on a monitor. (D) Camera panel (top middle panel; C): camera operations, FPS and stream attached predictors. (E) The arena settings and configuration (cog icon on the top left side of the UI). (F) Creating experiment panel

**Figure S5:**
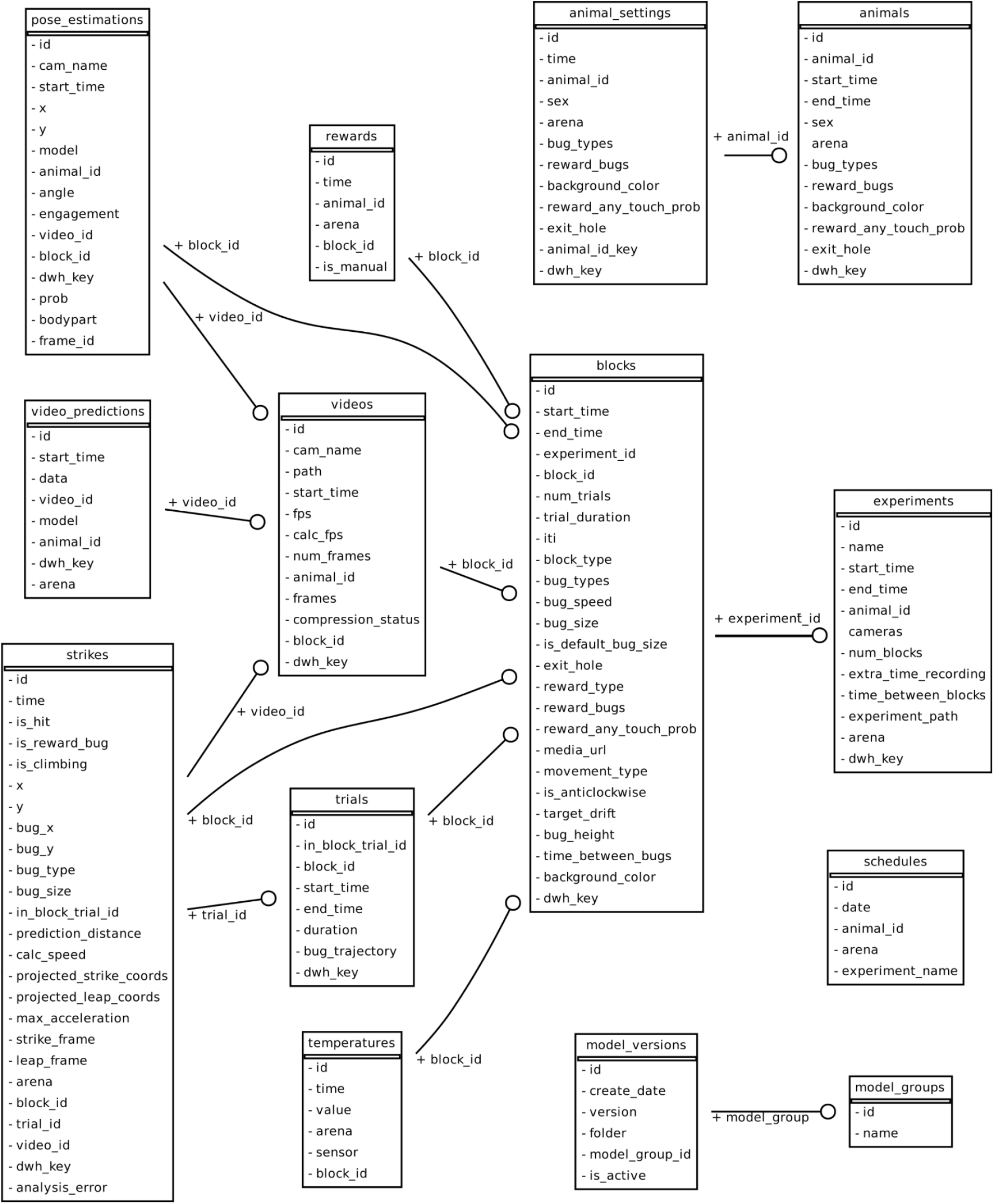
Database Schemas. A visualization of the PreyTouch database tables and their relations. Lines with circle markers indicate a relation between 2 tables, while the text below each line is the name of the foreign key in the source table.

